## Supplementary figures for "PSIP1/LEDGF reduces R-loops at transcription sites to maintain genome integrity": Supplementary Information.pdf

Supplementary file includes

Supplementary Figures 1 to 6

Supplementary Data 1 and 2

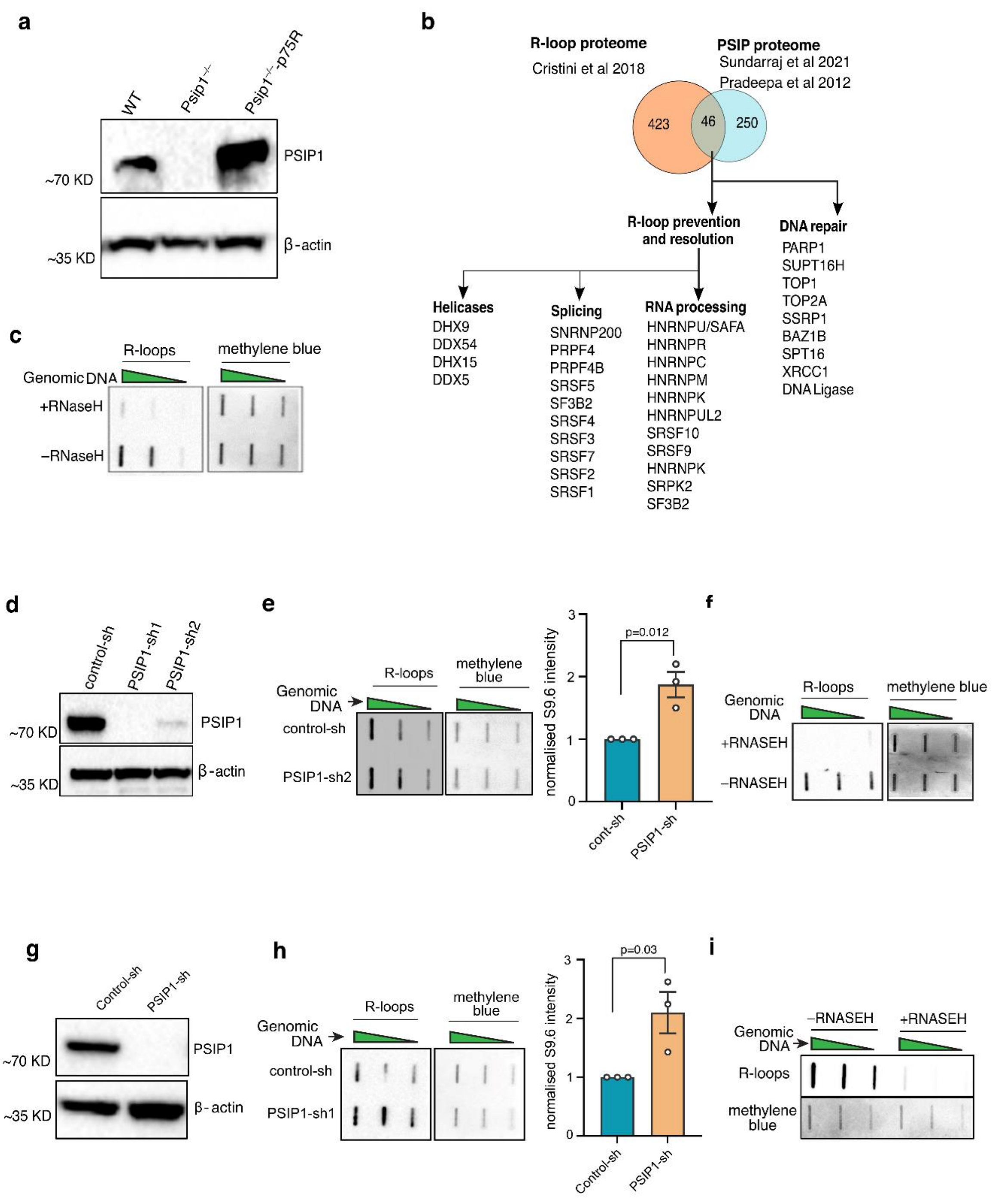


**Supplementary Figure 1.** ***PSIP depletion leads to increased R-loop levels.*** ***a****) Western blot image showing the levels of PSIP1 in wild type, PSIP knockout, and PSIP1 knockout re-expressed with P75 MEFs. β-actin was used as a loading control.* ***b****) The list of proteins that overlap between the R-loop and the PSIP1 interactors is shown, along with their functions related to R-loop resolution and DNA repair.* ***c****) Slot blot for R-loop using S9.6 antibody for genomic DNA extracted from Psip1 knock out (Psip1–/–) MEFs and treated with RNASEH. The same membrane was treated with methylene blue for the loading control.* ***d****) Western*

*blotting of RWPE-1 cells transduced with lentiviral shRNAs targeting PSIP1 (PSIP1-sh) with PSIP1/p75 antibody. Non-targeting control shRNA (control-sh) was used as control.* ***e****) Slot blot of S9.6 antibody for genomic DNA extracted from PSIP1 knockdown (PSIP1-sh2) and control knockdown (control-sh) RWPE-1 cells. The same membrane was stained with methylene blue to detect DNA as loading control. The intensity of R-loops in control and PSIP1 KD RWPE-1 cells in slot blot was normalised with DNA content and the relative normalised intensity of S9.6 as mean ± SEM has been plotted (right) (n=3 independent experiments; p values by two tailed unpaired t-test).* ***f****) Slot blot for R-loop using S9.6 antibody for genomic DNA extracted from PSIP1 KD RWPE-1 cells and treated with RNASEH. The same membrane was treated with methylene blue for the loading control.* ***g****) Western blot image showing the levels of PSIP1 after transfection with either control-shRNA or PSIP1-shRNA in HEK293T cells. β-actin is used as loading control.* ***h****) Similar to* ***e*** *but in HEK293T cells. The relative normalised intensity of S9.6 has been plotted as mean ± SEM (right) (n=3 independent experiments; p values by two tailed unpaired t-test)* ***i****) Slot blot for R-loop using S9.6 antibody for genomic DNA extracted from PSIP1 KD HEK293T cells and treated with RNASEH. The same membrane was treated with methylene blue for the loading control.*

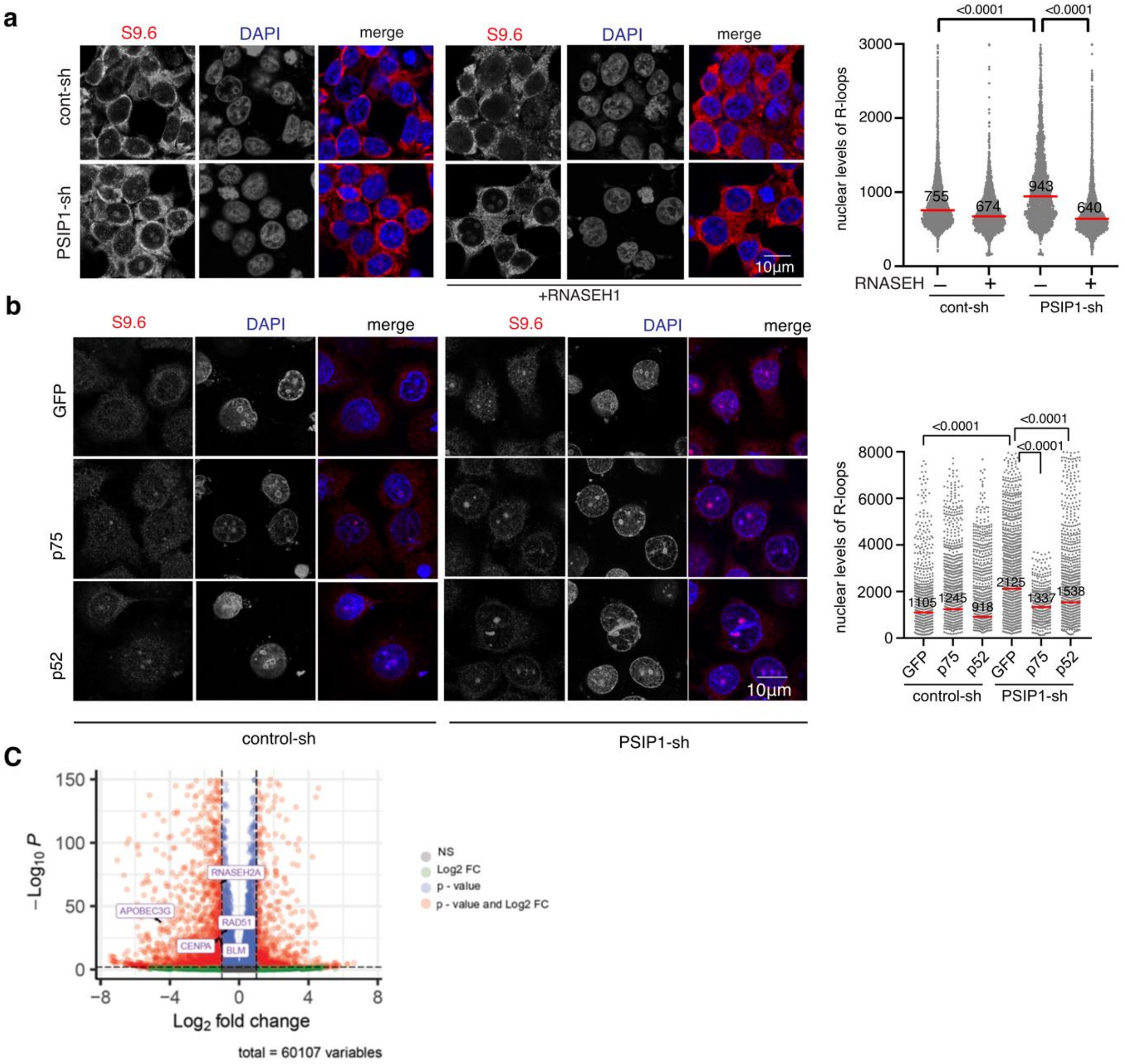


**Supplementary Figure 2.** ***PSIP depletion leads to increased R-loop levels***. ***a***) *Representative images and dot plot of S9.6 immunofluorescence in wild type and PSIP1-KD HEK293T cells either with or without RNASEH overexpression (representative images; n>3000 nuclei over three independent experiments; median values have been indicated with red line; p values obtained by two-tailed Mann-Whiney test).* ***b***) *Immunofluorescence images of S9.6 after transfecting with either GFP or p75 or p52 plasmids in control and PSIP1 KD cells. The S9.6 signal was also quantified and plotted as dot plot (n>650 nuclei over three independent experiments; median values have been indicated with red line; p values obtained by two-tailed Mann-Whiney test).* ***c***) *RNA-seq data showing differentially expressed genes in PSIP1 knockdown RWPE-1 cells as compared to control cells. We analysed for the differential expression level of 186 genes (Supplementary Data 1) that are known to have a role in R-loop homeostasis. Out of these genes, significantly dysregulated upon PSIP1 depletion are highlighted.*

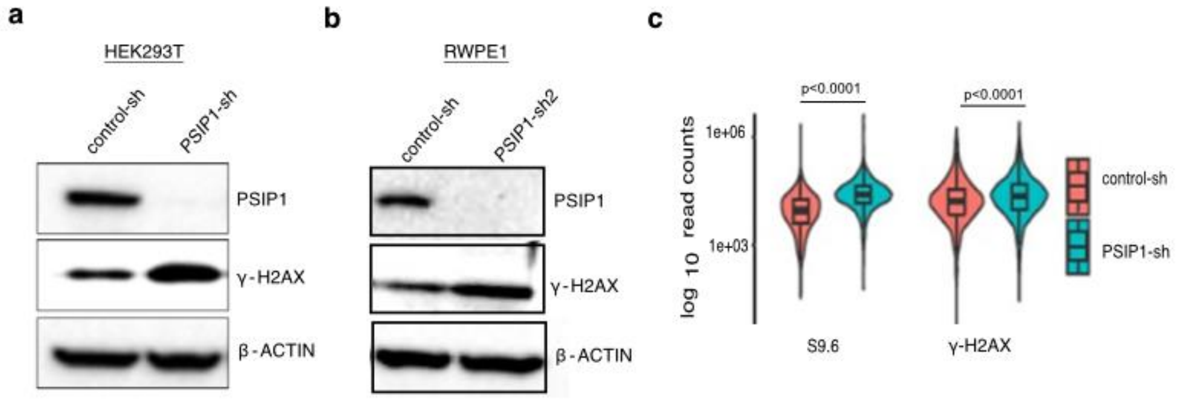


**Supplementary Figure 3.** ***a**) Western blot image showing the levels of PSIP1 and γ-H2AX in HEK293T cells transduced with either control shRNA or PSIP1 shRNA. β-actin was used as the loading control. **b**) Western blot image showing the levels of PSIP1 and γ-H2AX in RWPE-1 cells transduced with shRNA-2 targeting PSIP1. **c**) Violin plot showing E. coli normalised CUT&Tag (log10 CPM) for S9.6 and γ-H2AX antibodies at S9.6 peaks gained in KD. *** p<0.001 in Dunn test with Bonferroni correction.*

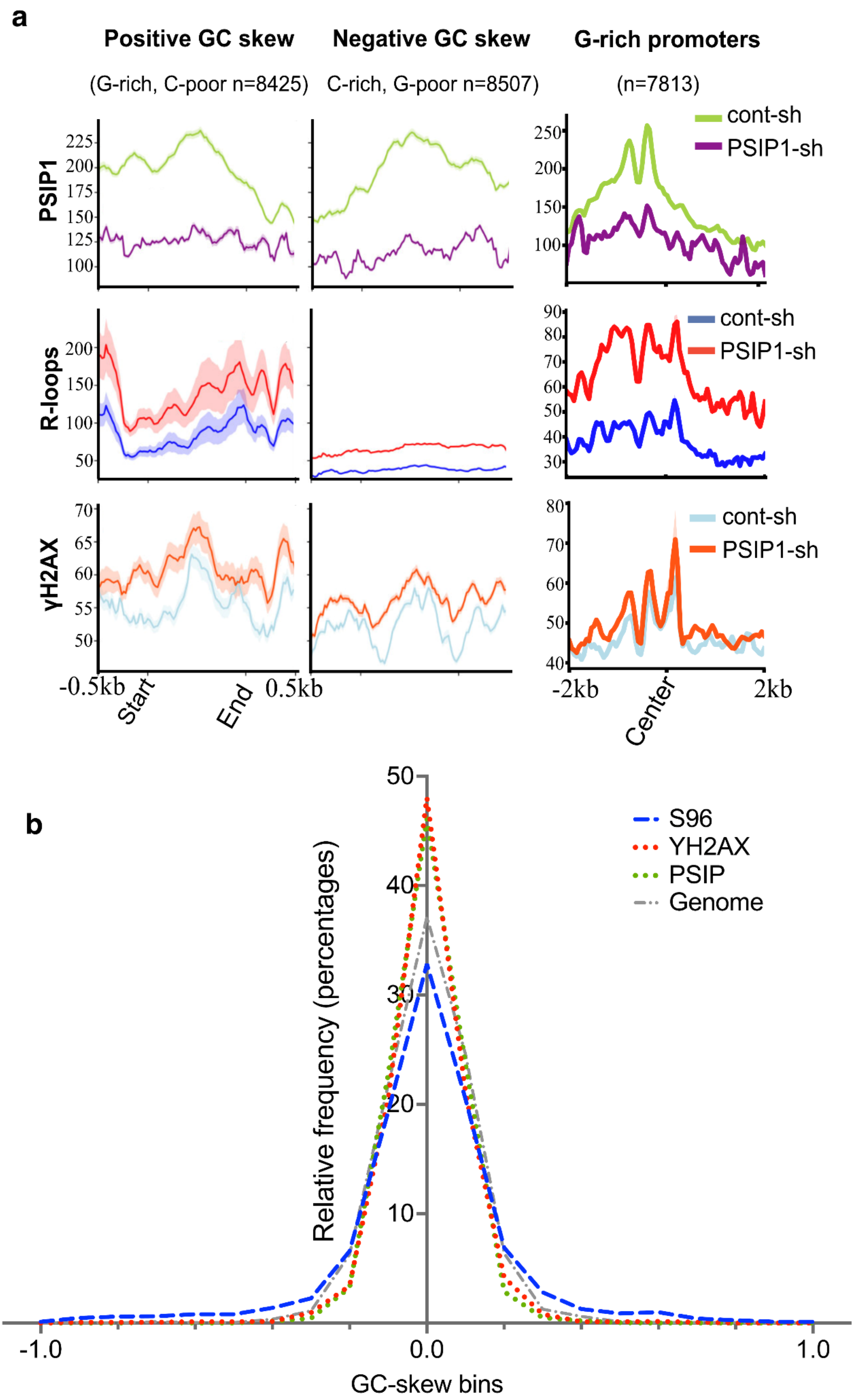
a
Positive GC skew
(G-rich, C-poor n=8425)
Negative GC skew
C-rich, G-poor n=8507)
G-rich promoters
(n=7813)
PSIP1
R-loops
γH2AX
cont-sh
PSIP1-sh
cont-sh
PSIP1-sh
cont-sh
PSIP1-sh
-0.5kb
Start
End
0.5kb
-2kb
Center
2kb
b
S96
YH2AX
PSIP
Genome
Relative frequency (percentages)
GC-skew bins
-1.0
0.0
1.0

**Supplementary Figure 4.** ***a****) CUT&Tag data showing the enrichment of PSIP1/p75 (top), R-loops (middle) and γ-H2AX (bottom) across positive GC skew, negative GC skew regions along with G rich promoters, in control (cont-sh) and PSIP1 knockdown (PSIP1-sh) RWPE-1 cells.* ***b****) GC skew calculated for the PSIP1 peaks found in the wild-type cells, S9.6 peaks, and γ-H2AX peaks found in the PSIP1 depleted cells. Randomly selected regions were used as the control representing the genome.*

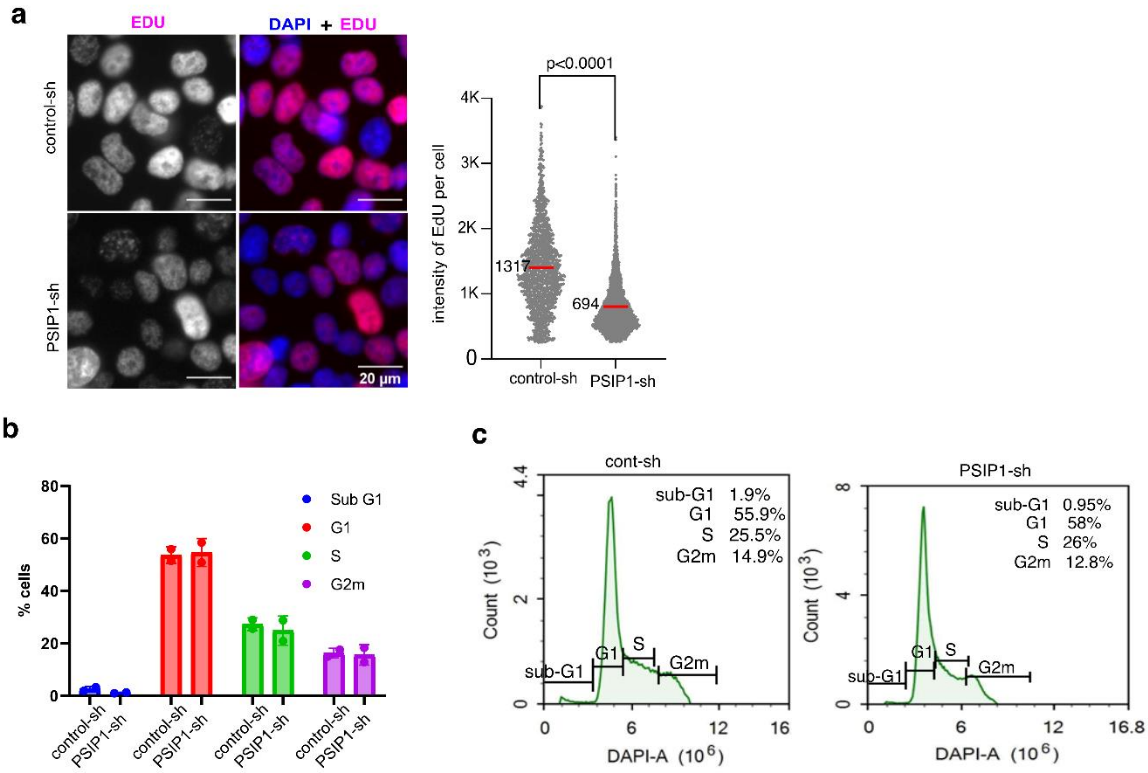


**Supplementary Figure 5.** ***a****) IF image and dot plot showing the incorporation of EdU in PSIP1-KD and control RWPE-1 cells (representative images; n>1600 nuclei over three independent experiments; median values have been indicated with red line; p values obtained by two-tailed Mann-Whiney test).* ***b****) Graph showing the percentage of cells in different cell cycle phases in control and PSIP1 KD condition as assessed by flow cytometry.* ***c****) Representative histogram of control (left) and PSIP1 KD cells (right) with their distribution across the different cell cycle phases.*

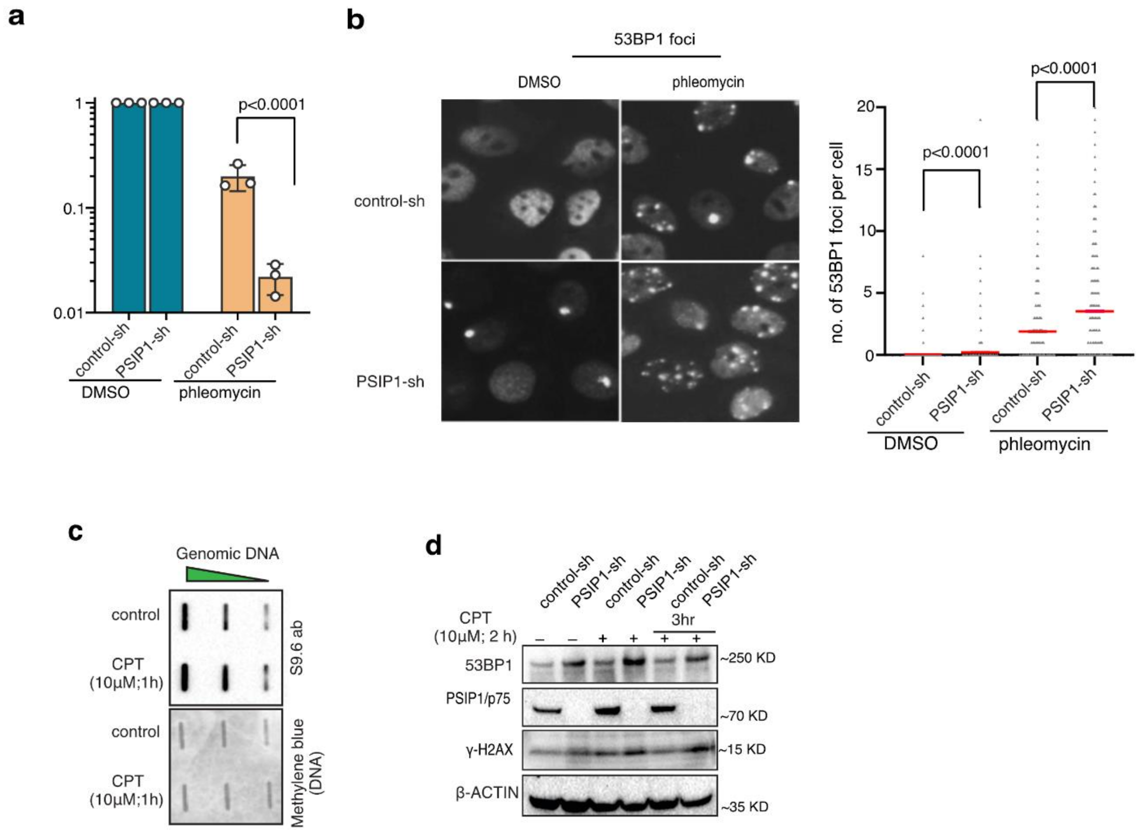


**Supplementary Figure 6.** ***a****) Cell survival fraction estimated by clonogenic assay with phleomycin treatment (0.1µg/ml) in RWPE-1 cells. (mean ± SD is plotted; n=three independent experiments; p values obtained by two-way ANOVA followed by Šídák's multiple comparisons test).* ***b****) Representative microscopic image of immunofluorescence showing 53BP1 foci in control and phleomycin-treated (1µg/ml) RWPE-1 cells. The number of foci per cell is shown as a dot plot (right) (n>6600 nuclei over three independent experiments; red line indicates the median; p values from two-tailed Mann-Whitney test).* ***c****) Slot blot for R-loop using S9.6 antibody for genomic DNA extracted from HEK293T cells treated with CPT (10 µM; 1 hrs). The same membrane was stained with methylene blue to detect DNA as a loading control.* ***d****) Western blot for γ-H2AX and 53BP1 in control and PSIP1-KD HEK293T cells treated with CPT (10µM; 2hrs) followed by repair for the indicated time. β-actin served as a loading control.*
